# Evaluating the effects of data visualisation techniques on interpretation and preference of feedback reports

**DOI:** 10.1101/415414

**Authors:** Harvey Koh, Arul Earnest, Ian D. Davis, Erwin Loh, Sue M Evans

**Author notes:** Corresponding author: Prof. Susan Evans., Department of Epidemiology and Preventive Medicine, Monash University, Level 2, 553 St Kilda Road, Melbourne 3004, AUSTRALIA.

## Abstract

**Objective:** To evaluate the different methods of data visualisation and how it affects preference and data interpretation.

**Design:** A cross-sectional survey, assessing interpretation and preference for four methods of data presentation, was distributed to participants.

**Setting:** Melbourne, Victoria

**Participants:** Members of Prostate Cancer Outcome Registry-Victoria (PCOR-Vic) and senior hospital staff in three metropolitan Victorian hospitals.

**Interventions:** Different methods of data visualisation. Mainly, funnel plots, league charts, risk adjusted sequential probability ratio test (RASPRT) charts and dashboard.

**Main Outcome Measure:** Interpretation scores assessed capacity by participants to identify outliers and poor performers. Preference was based on a 9-point Likert-scale (0 – 9).

**Results:** In total, 113 participants responded to the online survey (16/58 urologists and 97/297 senior hospital staff, response rate 32%). Respondents reported that funnel plots were easier to interpret compared to league charts (mean interpretability score difference of 28% (95% CI: 19.2% - 37.0%, P<0.0001). Predictors of worse interpretability of charts in the adjusted model were being a hospital executive compared to a urologist (coefficient= −2.50, 95% CI = −3.82, - 1.18, P<0.01) and having no statistical training compared to those with statistical training (coefficient = −1.71, 95% CI=-2.85, −0.58, P=0.003). Participants preferred funnel plots and dashboards compared to league charts and RASPRT charts (median score 7/9 vs 5/9), and preferred charts which were traffic-light coloured versus greyscale charts (43/60 (71.6%) vs 17/60 (28.3%)).

**Conclusion:** When developing reports for clinicians and hospitals, consideration should be given to preference of end-users and ability of groups to interpret the graphs.

## Introduction

Clinical quality registries (CQRs) are data management systems designed to implement quality assurance and quality control processes to provide consistent feedback to clinicians and treatment centres. CQRs are responsible for the secure collection of a defined dataset from patients who undergo a specific medical intervention, are diagnosed with a certain disease or use a healthcare resource.[1, 2] The collected datasets are then developed into multiple quality indicators through a rigorous selection process.

Quality indicators can be defined as “*quantitative measures that can be used to monitor and evaluate the quality of important governance, management, clinical, and support functions that affect patient outcomes”*.[3] Quality indicators collected and reported by CQRs usually monitor dimensions of quality such as whether care delivery is safe, efficient, equitable, effective, patient-centred and timely.[4] Data on the selected quality indicators are then collated and analysed with results often being presented using a graphical/ tabular format. Findings are incorporated into periodic performance reports comparing clinicians or treatment centres with their de-identified peers, and these are generally distributed to clinicians and senior hospital staff in treatment centres.[5]

CQRs have been proven to be effective as a clinical intervention tool, reducing adverse outcomes and improving quality of care through benchmarking processes and outcomes of care, and identifying areas of concern for action.[6] Established in 1994, the Riks-Stroke Registry in Sweden has demonstrated a reduction in adverse outcomes from strokes which has been attributable in part to the annual feedback provided to physicians and hospitals via quality indicator reports.[7]

The implementation of CQRs has led to improvements in processes and outcomes of care.[6] A conservative estimate of the benefit-to-cost ratio of highly developed CQRs post-implementation was recently estimated to range from a ratio for 2:1 to 7:1.[8] For the period 2009 to 2013, the Prostate Cancer Outcomes Registry – Victoria (PCOR-Vic), achieved a 2:1 benefit-to-cost ratio with capture of just 11% patients diagnosed in Australia (number of participating hospitals nationwide). Theoretically, with full national coverage, the estimated extrapolated benefit-to-cost ratio of the Prostate Cancer Outcomes Registry could be as high as 5:1.[8]

It is unclear whether the complex information presented by CQRs is achieving its optimal impact. Studies have shown that healthcare professionals may not be able to interpret data presented to them in visual formats.[9-11] Therefore, there is merit in understanding the interaction between data presentation and interpretation. Data presentation seeks to find the most efficient and effective way of translating raw data into presentable, interpretable and preferably, visually appealing visual aids that can aid a user in eventual decision making.[12] In the context of healthcare, these graphical methods are used by CQRs in clinical reports to inform healthcare providers on quality of care.

Quality Indicator reports developed by CQRs are designed to be easily interpretable and are intended to guide good decision-making and improvement in quality of care. The aim of this study was therefore to evaluate the different methods of data visualisation and how it affects preference and data interpretation in two different groups: urologists and senior hospital staff. In addition, we sought to explore preferences for visual aids (colour codes) in these two groups. We hypothesised that there would be difference in capacity to interpret charts based on profession, type of chart, level of statistical education, self-rated statistical knowledge and self-rated ease of interpretation.

## Methods

### Study design

A cross-sectional survey with a structured questionnaire was designed to be self-administered via the online survey tool, Qualtrics.^®^ Four different data presentation methods were tested (Fig 1): (1) a league chart, which uses the traditional bar chart to rank the performance of healthcare providers from highest to lowest, with each bar representing a provider. (2) a funnel plot, a traditional control chart with decreasing confidence intervals as the case number increases. (3) A risk-adjusted sequential probability ratio test (RASPRT) chart; a time-series control chart; and (4) a dashboard, which provides a summary of progress over the previous four reporting periods using simple coloured dots to indicate whether performance was better than, on par with or worse than the performance of peers. The dashboard was inspired by research undertaken by Guthrie et al. which aimed to summarise data from multiple quality indicators.[13] The progression bar is based on Tufte’s Sparkline, a proposed way of visualising progress through time.[12]

**Figure 1.**
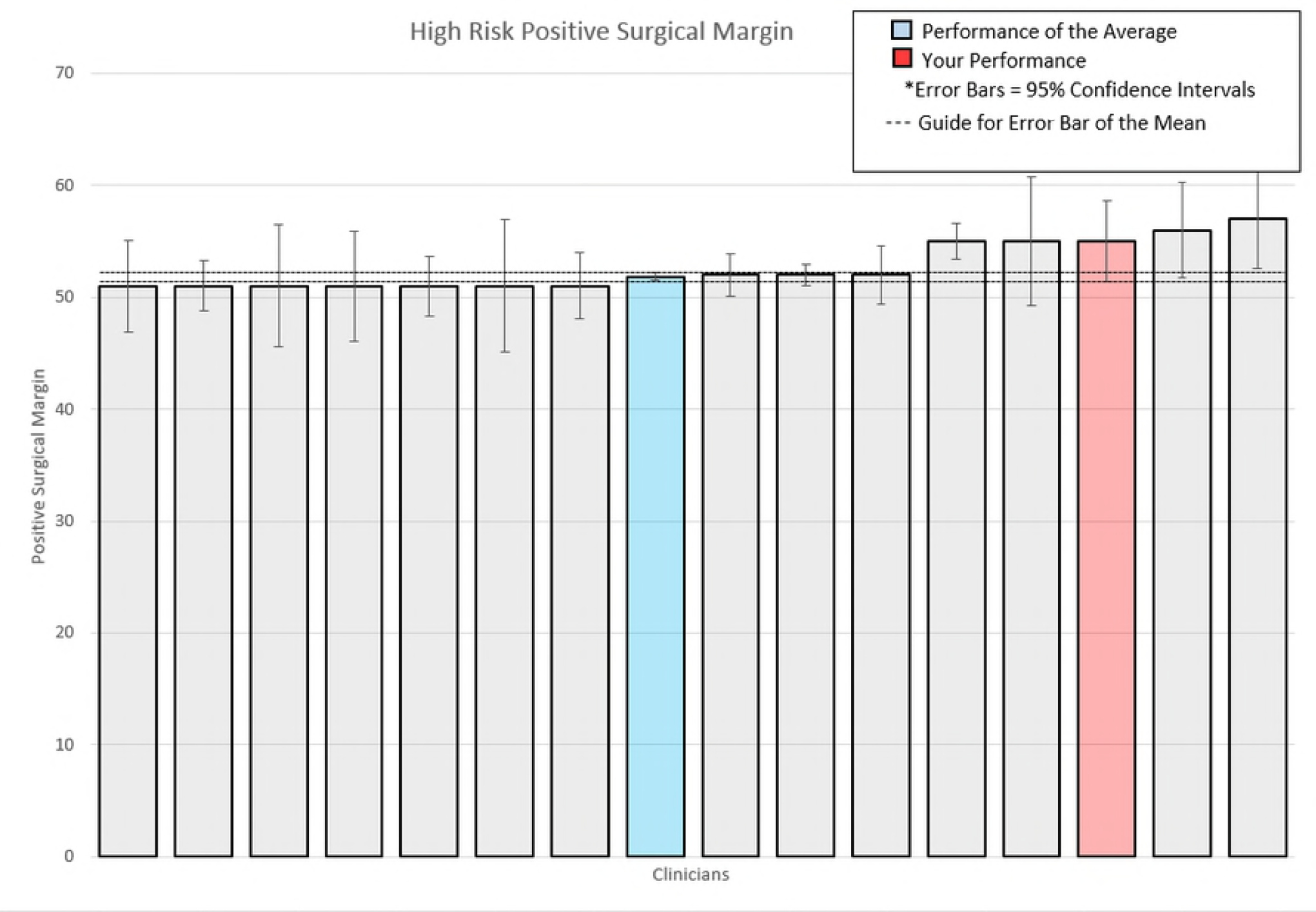
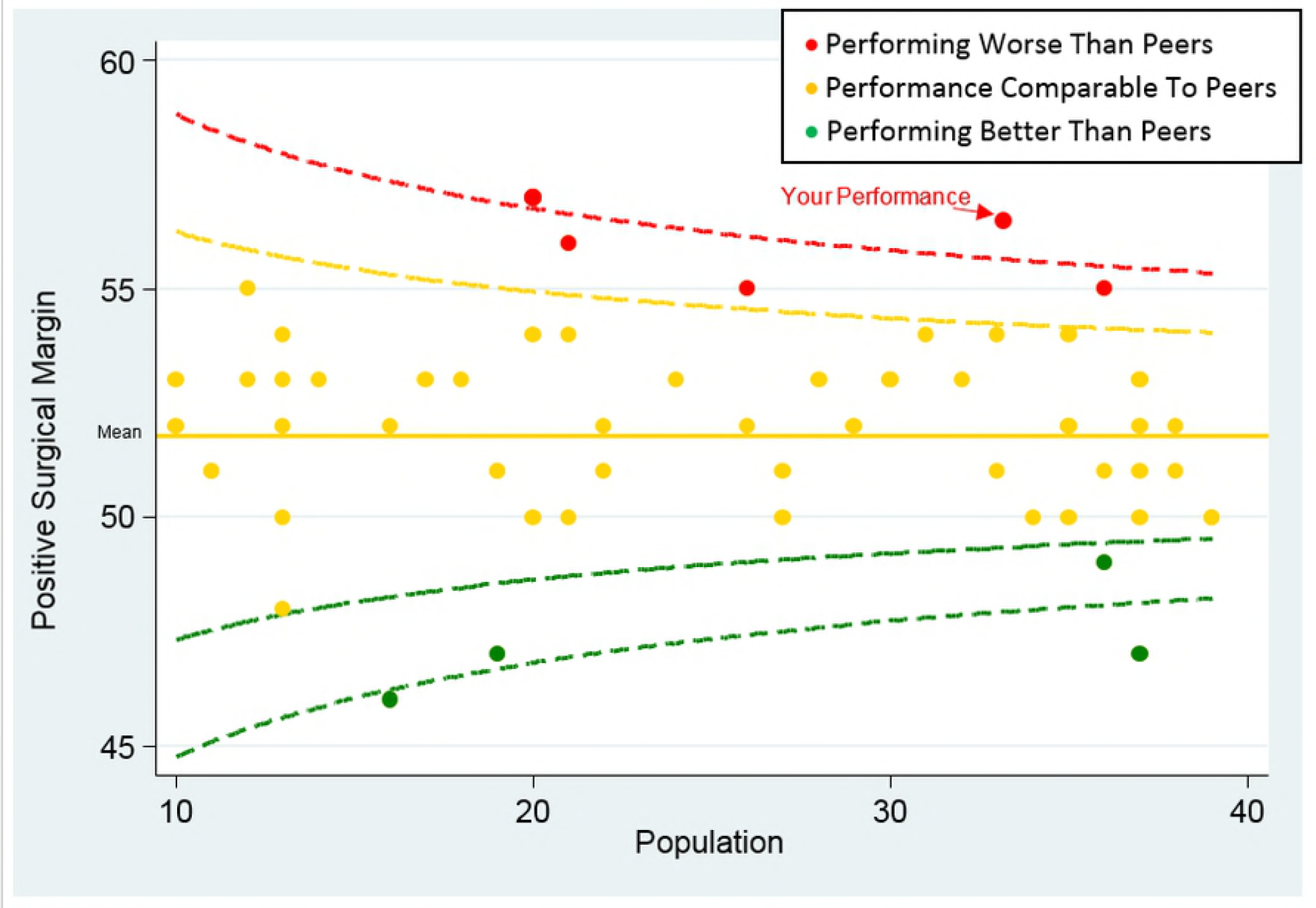
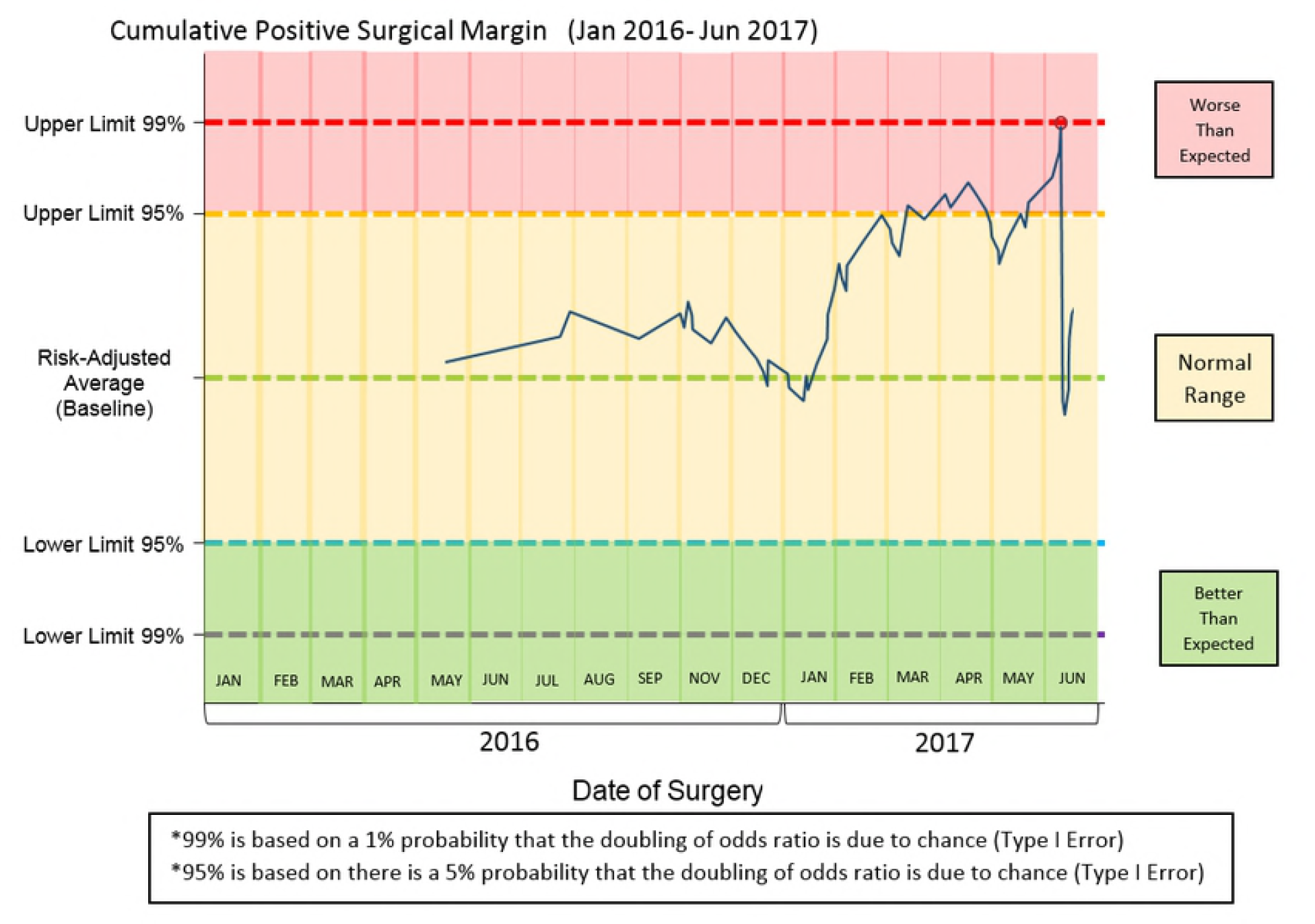
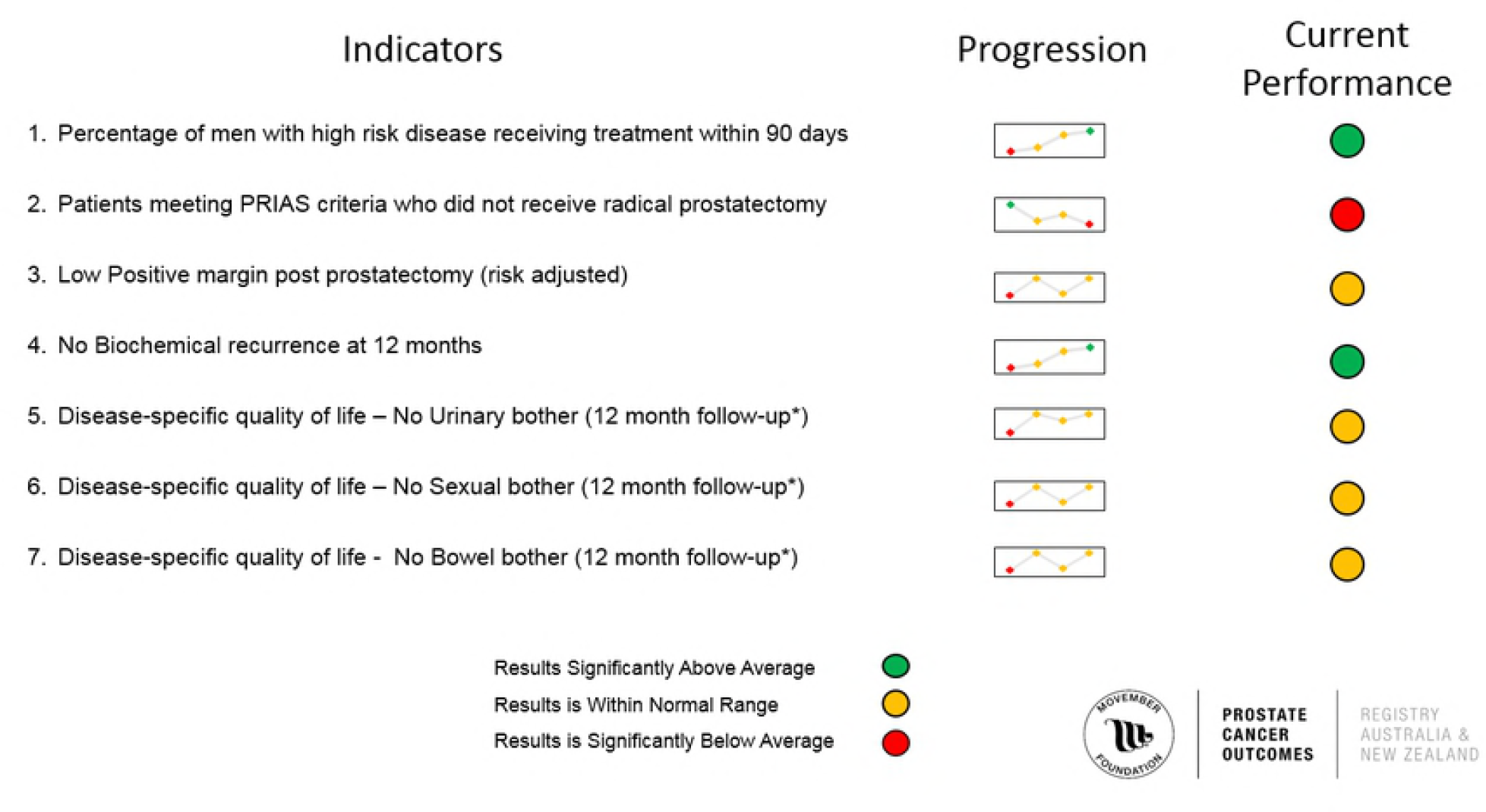
The types of data visualisation techniques used in this study. For all figures, red represents a performance worse than 95% of peers, amber represents within normal range and green represents performance that is better than 95% of peers. (A) League charts. (B) Funnel Plot. (C) Risk-Adjusted Sequential Probability Ratio Test. A chart which uses statistical process control to benchmark clinicians which is implemented in the survey. Figure 1D. Business Analytics Dashboard. Dashboard which allowed a summary of all quality indicators provided in the report.

The survey consisted of the following three components:

1. Component 1 assessed whether respondents could correctly answer questions relating to the data (“interpretability score”). It was calculated by assessing the extent to which respondents correctly answered ten questions, four focusing on funnel plots, two on league charts and four on RASPRT charts. These questions asked participants to denote whether a hospital was statistically different from the mean and if it was performing poorly compared to peers. Before each graph was shown, a short introduction of each graph as well as a ‘how to interpret guide’ was given to participants (supplementary document).

2. Component 2 assessed participant’s preference (“preference score”) for the different data displays. It was calculated by asking participants to answer on a 9-point Likert Scale how easy (easiness of interpretation score) each of the four types of graphical displays were to interpret (league charts, funnel plots, RASPRT and dashboard); and (2) the extent to which they liked the way the data was presented. A third additional question was asked for additional comments on how to improve the visuals.

3. Component 3 assessed respondents’ views on colour scheme for funnel plots (greyscale vs coloured funnel plots).

Demographic questions assessed participants’ statistical education. Participants were asked if they had formal (academic qualification level or equivalent), informal (education outside of an education institution) or no statistical training and to self-rate their confidence in understanding basic statistical concepts using a 9-point Likert scale.

#### Participants

Urologists and senior hospital staff were invited to participate in the survey. Urologists who contributed to the PCOR-Vic and had received at least one prior clinician quality indicator report were invited via electronic mail (email) with a link to the survey sent by the PCOR-Vic coordinator. Senior hospital staff included medical, nursing and allied health personnel from three metropolitan hospitals in Melbourne, Victoria. Senior hospital staff have a high level responsibility for clinical quality and safety in the acute hospitals contributing to the study. They were invited via email by the executive sponsor at each hospital. Known strategies to maximise recruitment were adopted, including sending personalised letters of invitation and two reminders.[14]

#### Statistical analysis

Descriptive statistics were used to assess the characteristics of the study participants. Paired t-test was used to compare the difference in mean percentage of questions answered correctly between the two cross sectional charts (funnel plots and league charts). To score participant preference, a median and inter-quartile range was calculated for each chart and categorised as follows: a median of 0-3 was defined as unfavourable, 4 – 6 was considered neutral, and 7-9 was considered favourable. Univariate and multivariate linear regression was used to analyse the relationship between interpretation score (questions answered correctly out of ten) against three predictors; (1) statistical education, (2) self-rated confidence in understanding basic statistical concepts and (3) the mean score of easiness of interpretation based on three charts (funnel plot, league charts and RASPRT). A chi-squared test was performed to test the questions comparing the preferences of colour coding between groups.

#### Sample size calculation

Based on the assumption that the 2-sided level of significance was set at 5% and that the standard deviation of the score was 2, a total sample size of 128 was required for a power of 80% to detect at least a clinically significant difference of 1 score between urologists and senior hospital staff. After the projected non-response rate of 60%, we calculated a final target sample size of 320. Data were analysed using STATA 13. Level of significance was set at 5%.

The ethical conduct of this multi-site study was approved by the Alfred Health Human Research Ethics Committee (HREC/17/Alfred/104) with governance approval provided by each contributing hospital.

## Results

Characteristics of respondents are described in Table 1. A total of 355 surveys were distributed and 113 responses were received (16/58 urologists (28%) and 97/297 (33%) senior hospital staff) providing a response rate of 32%. Most respondents stated they had official statistical education (65/112, 57.5%) and Intermediate (4-6) confidence in understanding basic statistical concepts (64/113, 56.6%)

**Table 1.**
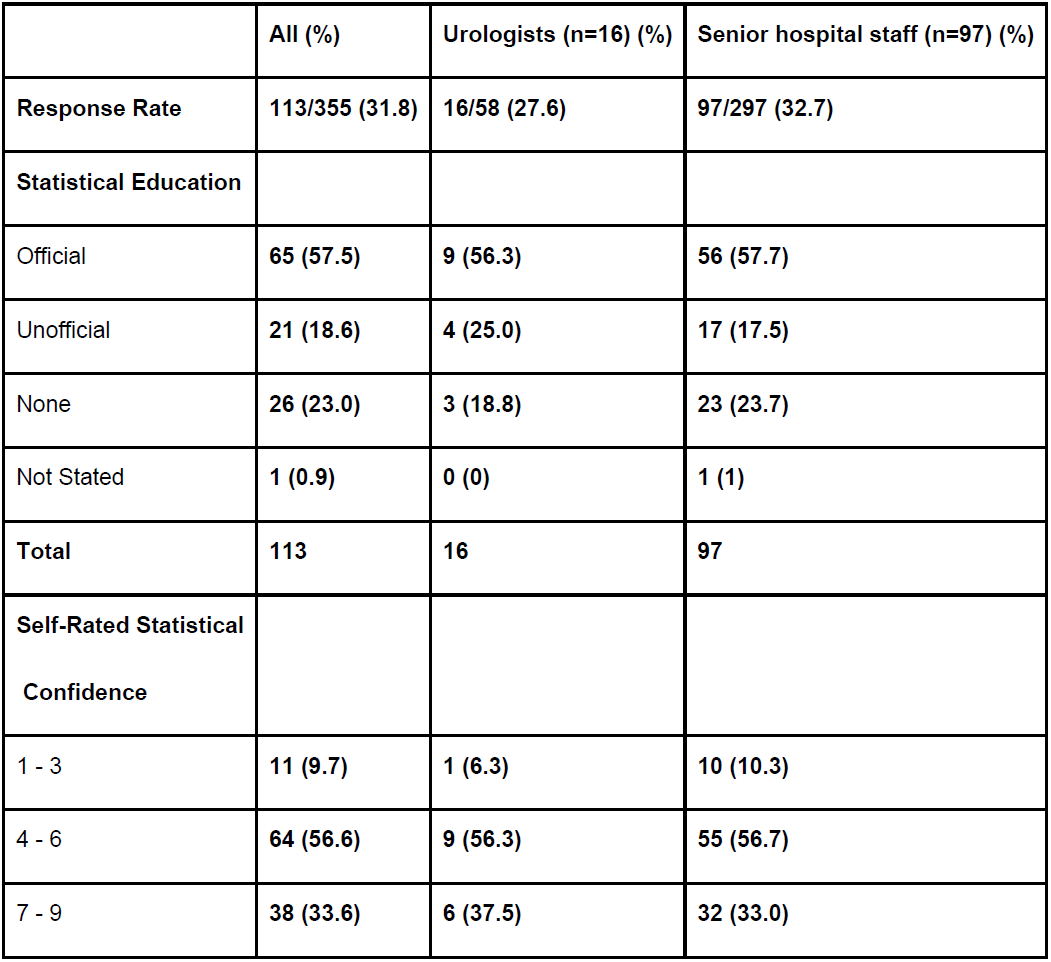
Characteristics of Participants.

### Interpretability

Overall, participants had an overall mean interpretation score of 43.3% (Table 2). Funnel plots scored significantly higher than league charts (69.7% vs 41.6%, p<0.001). RASPT charts had an interpretability score of 48.7%.

**Table 2.** Summary of Interpretation Scores of Different Charts in Percentage.

|  | Percentage<sup>#</sup> (%) | Standard Deviation | Significance<sup>*</sup> |
| --- | --- | --- | --- |
| **Cross-Sectional Charts** |  |  |  |
| Funnel Plot Mean Score | 69.7 | 28.7 |  |
| League Chart Mean Score | 41.6 | 44.1 | $P<0.001$ |
| **Time-Series Charts** |  |  |  |
| RASPT Mean Score | 48.7 | 41.1 |  |
| **Total** | 53.3 |  |  |

<sup>\*</sup>Paired T test used
<sup>#</sup>Mean percentage of questions scored correct for each graph.

Table 3 outlines the results of the univariate and multivariate analyses. Univariate linear regression demonstrated that senior hospital staff were significantly less likely than urologists to correctly interpret the graphs, on average, answering 2.5 questions less correctly compared with urologists (coefficient= −2.54, 95% CI = −3.90, −1.18, P<0.01). The unadjusted model demonstrated that a participant’s statistical education significantly impacted their interpretability score (coefficient = −1.77, 95% CI = −2.97, −0.57, P<0.004). However, participants’ self-rated confidence in statistical knowledge was not a significant predictor of having a high interpretability score.

**Table 3.**
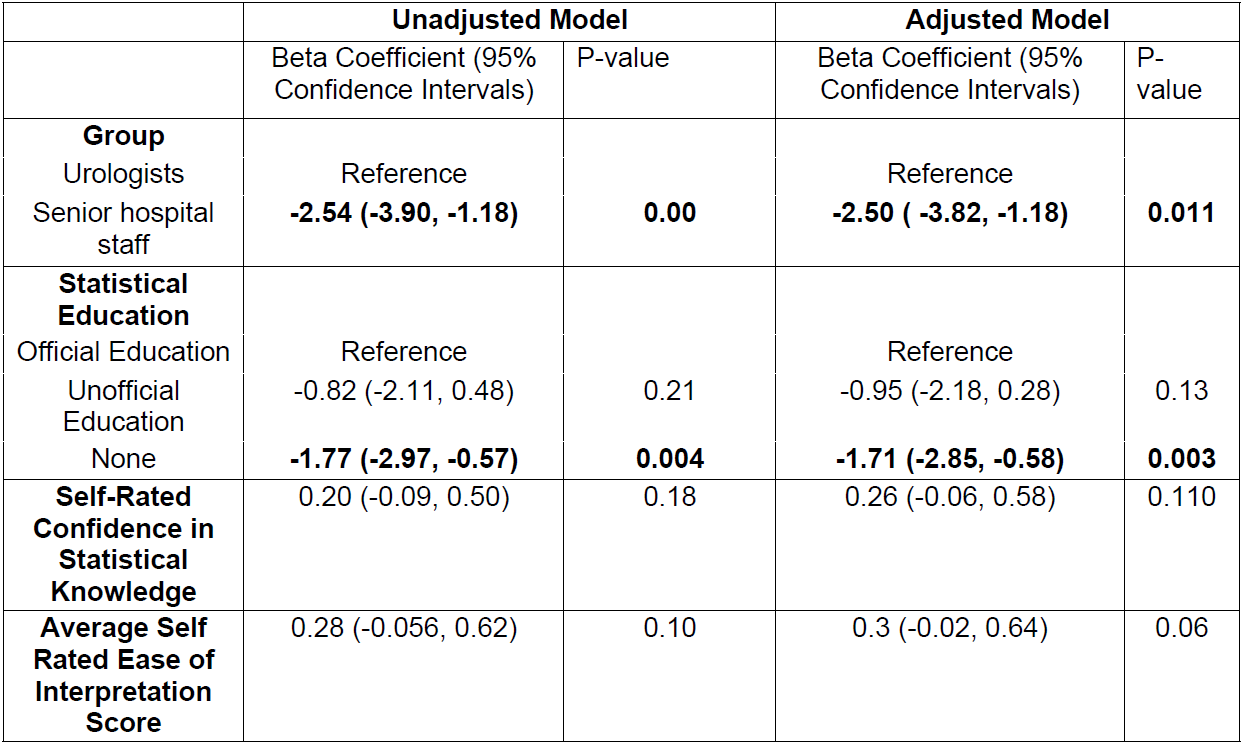
Predictors of Interpretation measured based on composite score of Interpretation.

After adjustment using stepwise multivariate regression, senior hospital staff scored significantly lower than urologists, answering 2.5 questions less correctly than urologists; a statistically significant result (coefficient= −2.5, 95% CI = −3.82, −1.18, P<0.001). Statistical education had a significant impact on interpretation scores, with participants with no statistical education answering 1.7 questions less correctly than participants with a statistical education (coefficient = −1.71, 95% CI=-2.85, −0.58, P=0.003). Self-rated confidence in statistical knowledge and average ease of interpretation score did not impact the interpretation score.

### Preference

Table 4 provides a summary of preference scores recorded by urologists and senior hospital staff for each of the different graphs. Participants rated highly their preference for funnel plots and dashboards, with each scoring a median score of 7 out of 9. Participants were less likely to prefer league charts and RASPRT with a median score of 5 out of 9. Urologists rated league charts and RASPRT lower than senior hospital staff (median score of 3 and 3.5 vs 5).

**Table 4.** Summary of results for preference.

|  | Median Preference Score (Scored out of 9) |  |  |
| --- | --- | --- | --- |
|  | All | Urologists | Senior hospital staff |
| **Type of graphical display (n=71)** |  |  |  |
| Funnel Plot | **7** | **7** | **7** |
| League Charts | **5** | **3.5** | **5** |
| RASPRT | **5** | **3** | **6** |
| Dashboard | **7** | **6.5** | **7** |

### Colour preference

Participants preferred colour-coded graphs which highlighted participant’s performance using a traffic light colour scheme compared to two-colour charts (17/60 (28.3%) vs 43/60 (71.6%)). Senior hospital staff overwhelmingly preferred the colour coded plots, with 37/49 (76%) senior hospital staff choosing traffic light colour coding, while 5/11 (45%) urologists preferred the two-colour scheme funnel plots.

## Discussion

This explorative study was designed to assess clinicians’ and senior hospital staffs’ ability to interpret data visualisation methods used in CQR reports as well as gauging participants preferences and interest in new methods of presenting data for PCOR-VIC, specifically the RASPRT process control chart and a dashboard. The study found that (1) funnel plots were more readily interpretable when compared to league charts, (2) urologists were more likely to be able to interpret data presented in charts than senior hospital staff, and (3) funnel plots and dashboards were highly preferable when compared to league charts and RASPRT.

Funnel plots were easier for respondents to interpret than league charts, while RASPRT charts were considered difficult to interpret. When asked to interpret the league charts, instead of comparing themselves to the mean to identify outliers, participants were likely to answer based on their position on the chart. It is likely that the use of ranking in league charts was confusing participants. This was demonstrated by Schmidtke et al., in a study finding that participants were likely to identify “special-cause datum” or outliers when using funnel plots when compared to league charts. It is also important to note that this study was conducted in laypeople rather than hospital decision-makers.[15] Other studies have also reported that when participants are provided with league charts and funnel plots and asked to identify outliers, they tended to identify more outliers in league charts when compared to funnel plots.[11, 16] However, in a study by Marshall et al., participants self-reported that league charts were much more interpretable when compared to control charts.[11] The authors of the study explained that the widespread use of league charts in healthcare reporting in the UK[17] may have biased respondents to favour the familiar league charts rather than control charts.

Our results, which show high interpretation scores for funnel plots, may also be biased by our sampling population. While league charts are used in 80% of registry reports in Australia[18] and Sweden[19], PCOR-VIC has incorporated funnel plots into their CQR reports to ensure statistical accuracy.[20] The CQR report distributed by PCOR-VIC has 12 funnel plots and one league chart in its performance report. The discrepancy in results may be due to urologists and senior hospital staff being unfamiliar with league charts. A similar reason can be inferred for RASPRT, given that RASPRT was newly introduced in this study. It is highly likely that participants scored RASPRT charts lower when compared to funnel plots because they were unfamiliar to respondents. As shown in previous studies, training can be successfully delivered to assist in the interpretation and integration of statistical control charts such as RASPRT and funnel plots into practice.[21-24] Urologists were more capable of correctly interpreting data presented in CQR reports when compared to senior hospital staff, even after adjusting for statistical knowledge and confidence. This likely reflects the fact that the urologists who participated in the study receive clinician-level CQR reports on a six-monthly basis, while senior hospital staff are less likely to have the same level of exposure to these reports and, by default, to these types of graphs.

After adjusting for other predictors, statistical education was found to be significant in its effect on the interpretation score. Effective data visualisation seeks to present highly complex data using simple visual aids that can be readily interpretable by users.[12] Hospital clinical governance committees and Boards are responsible for ensuring that the hospital is providing safe and effective care. It is imperative that they correctly interpret the reports presented to them. A survey of health service Board members in Victoria revealed that only 26% were health practitioners, while most had a professional background in disciplines such as business, education and law. It is important that even without statistical education, people responsible for overseeing health services can interpret data presented in CQR reports. Therefore, further improvement to the current methods of data visualisation and the inclusion of a dashboard may assist in the effective interpretation of CQR reports.

League charts rank clinicians and healthcare units from best to worst performing, giving the perception that a particular unit may be doing worse than others, even when results are not statistically significant. A systematic review has found clinicians suffer tremendous emotional distress such as shame, guilt, fear, panic, shock and humiliation when faced with medical errors, in part due to the culture of medicine which does not tolerate mistakes.[25] Our finding that funnel plots were preferred to league charts, particularly for urologists, may reflect fear of being villainised by data presented in league charts.

Two new methods of presenting data were introduced in this study; the RASPRT chart and the dashboard. Dashboards are widely used in quality engineering and are defined as *“a tool used by management to clarify and assign accountability for the “critical few” key objectives, key indicators, and projects/tasks needed to steer an organization toward its mission statement”.*[26] With the increasing reporting requirements of Boards,[27] a dashboard may potentially reduce their burden by quickly identifying areas of interest. Conversely, RASPRT charts were found to be confusing, likely due to their “reset” function. The reset occurs when the data point hits the upper/lower 99% control limit, indicating a 1% chance in a doubling of odds ratio due to chance. At this point the performance line resets back to baseline, a feature that allows continuous monitoring of an individual’s performance. This however may be misinterpreted as an improvement in performance, rather than an alert that would be alarming to most.

It is theorised that the simplicity of the dashboards was key in participants favouring it over the RASPRT. This inference is supported by a systematic review by Hildon et al. which found that data had the most impact if presented in a simple format (“less is more”).[28] It is likely, however, that in time and with adequate information and training, participants would recognise the value in a statistically accurate time-series control chart. A study undertaken in Queensland on a similar time-series control chart found that these charts were more sensitive than traditional cross-sectional analyses in early and accurate detection of aberrance.[29]

This study suffers from several limitations. The response rate of 31.8% was modest and, because we avoided capturing demographic details to encourage people to respond, it meant that we were unable to understand how representative responders were to the population under investigation. Therefore, we cannot assume these results can be generalised to the broader population [30] Secondly, this novel survey tool was tested for face validity but other more formal validation testing is required to assess construct, criterion, content validity and reliability of the instrument. Thirdly, our study may not be powered to detect a difference which exists between urologists and health executives in regard to the relationship between statistical education and statistical confidence and interpretability of data. Also, the survey of senior hospital staff was conducted in only three public health services and only among surgeons, so it is unclear as to the generalisability of our findings to other locations, in other types of health settings (e.g. private sector, aged care rehabilitation) and among different specialty groups. Finally, we grouped all senior staff with responsibility for quality and safety into one group (i.e. ‘senior hospital staff’); making assumptions about individuals in this heterogeneous group should be undertaken with caution.

## Conclusions

Data displayed as funnel plots among urologists and senior hospital staff was superior in identifying outliers in cross-sectional data and was preferred among this cohort when compared to league charts. However, funnel plots display data at one point in time, and future research is required to understand how to best ensure that data displaying progress over time to healthcare professionals is both statistically accurate and easy to interpret. Looking to other industries, such as business and marketing, may provide innovative approaches for displaying complex information which may be applied in health.

## Acknowledgements

IDD is supported by an NHMRC Practitioner Fellowship (APP1102604). We are grateful to the clinicians and senior staff for contributing to the study, Professor Andrew Way for his support in distributing the survey and to Melissa Gillespie for administrative support.

